# Neural mechanisms of self-conformity

**DOI:** 10.64898/2026.07.22.740209

**Authors:** Juri Fujiwara, Philippe N. Tobler, Ken-Ichiro Tsutsui, Yoshikazu Ugawa, Satoshi Eifuku, Masato Taira

**Affiliations:** Department of Systems Neuroscience, Fukushima Medical University, Fukushima, 960-1295, Japan; Laboratory for Social and Neural Systems Research, University of Zurich, Zurich, 8091, Switzerland; Systems Neuroscience Laboratory, Graduate School of Life Sciences, Tohoku University, Sendai, 980-8577, Japan; Department of Cognitive Neurobiology, Graduate School of Medical and Dental Sciences, Tokyo Medical and Dental University, Tokyo, 113-8510, Japan; Department of Neurology, Fukushima Medical University, Fukushima, 960-1295, Japan; Department of Neurology, NHO Fukushima National Hospital, Fukushima, Japan

**Author notes:** Deceased (Masato Taira).

**Keywords:** conformity, social influence, self-consistency, decision-making, fMRI, prefrontal cortex

## Abstract

We conform not only to the opinions and behaviors of others but also to those of ourselves, suggesting that the motivation to appear consistent leads us to treat ourselves like others. However, the neural mechanisms of self-conformity remain unclear. We used a sequential facial attractiveness-rating task with ostensible reminders of participants’ own previous ratings to investigate self-conformity and compare it against group conformity. Rating data showed that participants on average conformed to both themselves and the group. While some individuals conformed excessively (over-conformity), others went against their previous behavior (anti-conformity), similarly in self and group conditions. Neurally, dorsomedial and ventrolateral prefrontal cortex activity correlated with both self- and group conformity. Moreover, self-versus group conformity engaged more medial versus lateral prefrontal regions, and over-versus anti-conformity more ventral versus dorsal regions. These results suggest that self- and group conformity rely on both common and dedicated neural mechanisms.

## Introduction

To facilitate our social life, we often adjust our decisions and judgments to conform to the groups we are part of. Social influence on our behavior has been extensively studied in social psychology, which proposed that we conform because we desire to behave according to social norms, to obtain social approval from others, and to maintain a favorable self-concept^1–3^. For a positive self-concept, we not only conform to others but also to our own judgments and behaviors because we want to appear consistent with ourselves. For example, in choice blindness experiments, people conform to their own purported behavior and provide reasons for their choices even if their actual behavior or choice differed^4^. Possible interpretations include that people tend to react in socially desirable ways or engage in a self-persuasion process when they face a choice endorsed by themselves, thus engaging in self-conformity^5–8^. At first sight, the basic mechanisms of social conformity and self-conformity seem similar, in that both involve conforming to public information. However, the source of the information differs between the two forms of conformity. While we use others as reference in social conformity, we need to refer to our own judgments and behaviors in self-conformity. Most previous studies of conformity have focused on the conformity to others and little is known about self-conformity.

A substantial body of evidence has characterized the neural basis of social conformity. In particular, activity in the rostral cingulate zone extending into dorsomedial prefrontal cortex (dmPFC) has been implicated in monitoring self-other conflict and predicting subsequent change in behavior^3,9–12^. Also, dorsolateral prefrontal cortex (dlPFC) and orbitofrontal cortex are involved in social influence associated with conformity^3,10,13^. However, these studies all focused on social conformity. Thus, it remains unclear whether the neural mechanisms of group conformity also underlie self-conformity. Moreover, the neural mechanisms underpinning conflict have rarely been separated from those underpinning conformity.

To investigate self-conformity and compare it to social conformity, we modified a facial attractiveness rating task used in previous studies on social conformity. In this task, participants rate a face before and after obtaining social information. People on average increase or decrease their second compared to the first rating if the rating of peers is higher or lower than the first rating^3,9,13^. Here, we increased the interval between the first and second rating and added one condition where participants observed their own previous rating between the first and second rating. If self-conformity arises from similar processes as social conformity, then we would expect common brain regions for the two conditions. Conversely, brain regions that mediate the self– other distinction (e.g., temporoparietal junction; TPJ or medial prefrontal regions) might differentiate between the two conditions.

## Results

### Behavioral results

#### Social and self-conformity

First, we measured individual conformity by calculating the change in ratings between pre-rating and main rating. Participants on average did not change their ratings in the true trials, whereas they decreased their ratings in false negative trials and increased them in false positive trials, conforming to both the group and their own previous rating (Fig. 2a). A two-way (two conformity types (group / self) x three reminder types (false negative / true / false positive)) repeated-measure ANOVA revealed a significant main effect of reminder types (F(2, 56) = 52.85, *p < 0*.*0001), but no significant main effect of conformity type and no interaction between the two. There were also no significant effects on reaction times*.

**Fig. 1.**
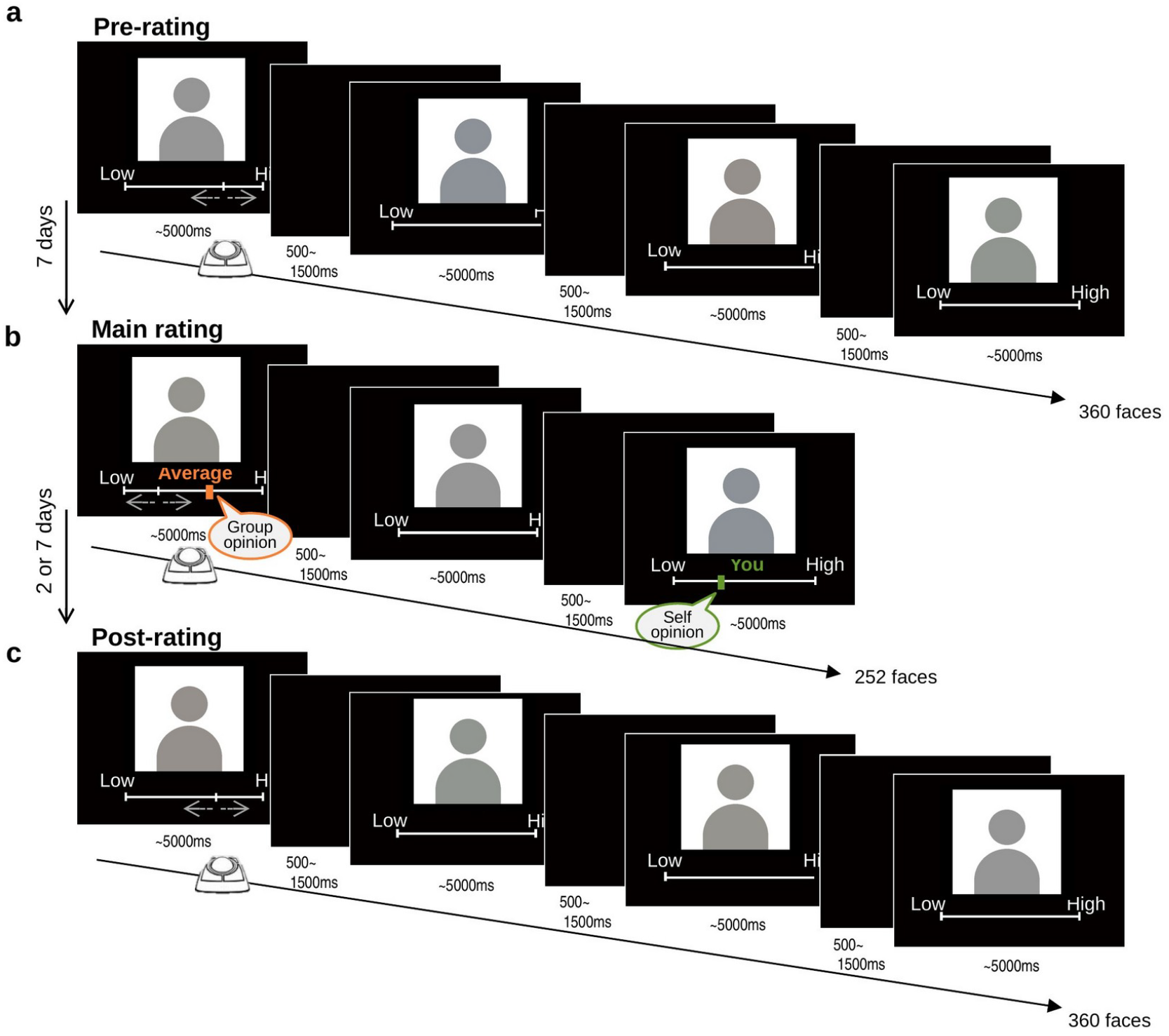
Experimental task. (a) Pre-rating task: participants rated the attractiveness of 360 female faces on a visual analogue scale. (b) Main rating task (one week later): each face was shown with the ostensible average rating of a previous group (“Average”), the participant’s ostensible own previous rating (“You”), or without additional information (“No reminder”); the reminder was either the true pre-rating (true trials) or shifted above or below it (false positive / false negative trials). (c) Post-rating task: faces were rated again without reminder to assess the temporal stability of conformity. In this figure the face stimuli are shown as schematic placeholder icons; the actual stimuli were photographs of real female faces (see Methods).

**Fig. 2.**
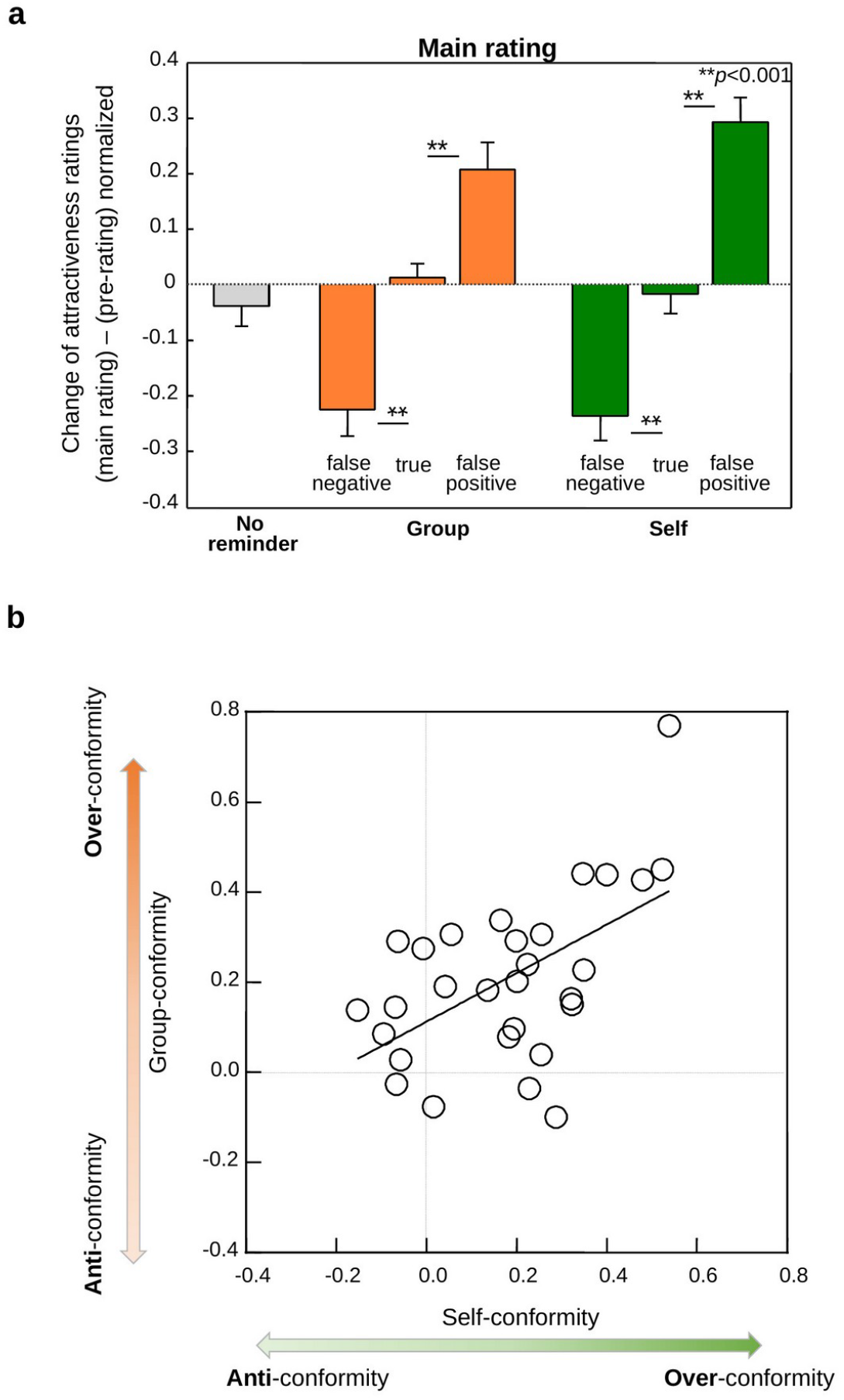
Behavior. (a) Mean rating change from the pre-rating to the main rating for the group and self conditions across false negative, true and false positive trials. (b) Relationship between the individual tendency to conform to the group and to oneself.

#### Over-conformity and anti-conformity

To characterize individual propensity to conform, we assessed each participant’s degree of conformity, pooling over false negative and false positive trials, and plotted group conformity and self-conformity separately (Fig. 2b). We found that the individual tendency to conform to a group of peers correlated positively with the tendency to conform to oneself (*r = 0*.*5, p < 0*.*005)*. Next, we split all trials in the main task into three groups, according to the sign and extent of conformity. First, trials where participants rated a face even more extremely than the indicated group or previous self rating (over-conformity), second, trials with a rating that had the same direction as the previous one but did not exceed the indicated rating (under- and iso-conformity, for readability labeled “under-conformity”) and third, trials with a rating in the opposite direction of the indicated rating (anti-conformity). Combining over-with under-conformity showed that participants conformed more often than they anti-conformed (59% versus 41%; t(28) = 6.02, p < 0.0001). Still, on average they anti-conformed more often than they either over- or under-conformed (F(2, 56) = 10.17, *p < 0*.*0005, Supplementary Fig. 2a)*.

#### Personality traits correlate with social and self-conformity

As another alternative approach to understanding conformity, we considered the Big Five model of personality traits. In particular, agreeableness and conscientiousness have been found to be related to prosocial behavior^14–16^ and may explain not only social but also self-conformity. In exploratory analyses we therefore assessed the correlation between the individual tendency to conform overall (averaging conformity with others and oneself) and personality inventory scores. We found a positive correlation between agreeableness and overall degree of conformity (*r = 0*.*4, p < 0*.*05; Supplementary Fig. 2*b). Moreover, stronger propensity to conform to oneself rather than the social group correlated with stronger conscientiousness (*r = 0*.*5, p < 0*.*01; Supplementary* Fig. 2c). Thus, relatively stable personality traits appear to be related to individual tendencies to temporarily conform.

#### Conformity is temporary

To examine the persistence of conformity over time, we administered a post-rating task in which participants rated the pre-rating images again, this time without reminders. Six participants were scanned two days after the main task, while the remaining participants were scanned one week after the task. Conformity in participants who performed the post-rating task two days after the main task did not significantly differ from that of participants with a seven-day interval between the two tasks. Importantly, in both cases, conformity observed in the main task returned to the initial ratings in the post-rating task (Supplementary Fig. 2d). Thus, in our task both self- and group conformity were similarly temporary.

### Neuroimaging results

#### Conflict-related regions

Following previous research^3^, we expected increased activity in conflict regions when the additional information (reminder) about the group differed from participants’ actual opinion. Thus, activation in conflict-detecting and monitoring regions should be increased during shifted compared to true trials. In the first step of identifying conflict-monitoring regions, we compared shifted to non-shifted reminder trials at the time of stimulus presentations. For the shifted trials, we pooled over false negative and positive trials (GLM1). We found that dmPFC, as well as dlPFC and precuneus, were more activated when participants viewed shifted compared to non-shifted reminders (Fig. 3a; whole-brain cluster-level FWE-corrected, p < 0.05, cluster-inducing threshold p < 0.001 uncorrected; this thresholding applied to all findings reported in the main text; Supplementary Table 1. No region showed stronger activity for non-shifted than shifted trials at the whole-brain cluster-level FWE-corrected threshold). Importantly, in these regions, conflict-related activity occurred in both social (group) and self conditions. For example, plotting the average effect size separately for the different trial types showed that dmPFC activity was similarly increased in both the social and the self conditions (Fig. 3b). Thus, a set of regions including dmPFC showed conflict-like activity that was common to situations where social and previous self information differed from current judgments.

**Fig. 3.**
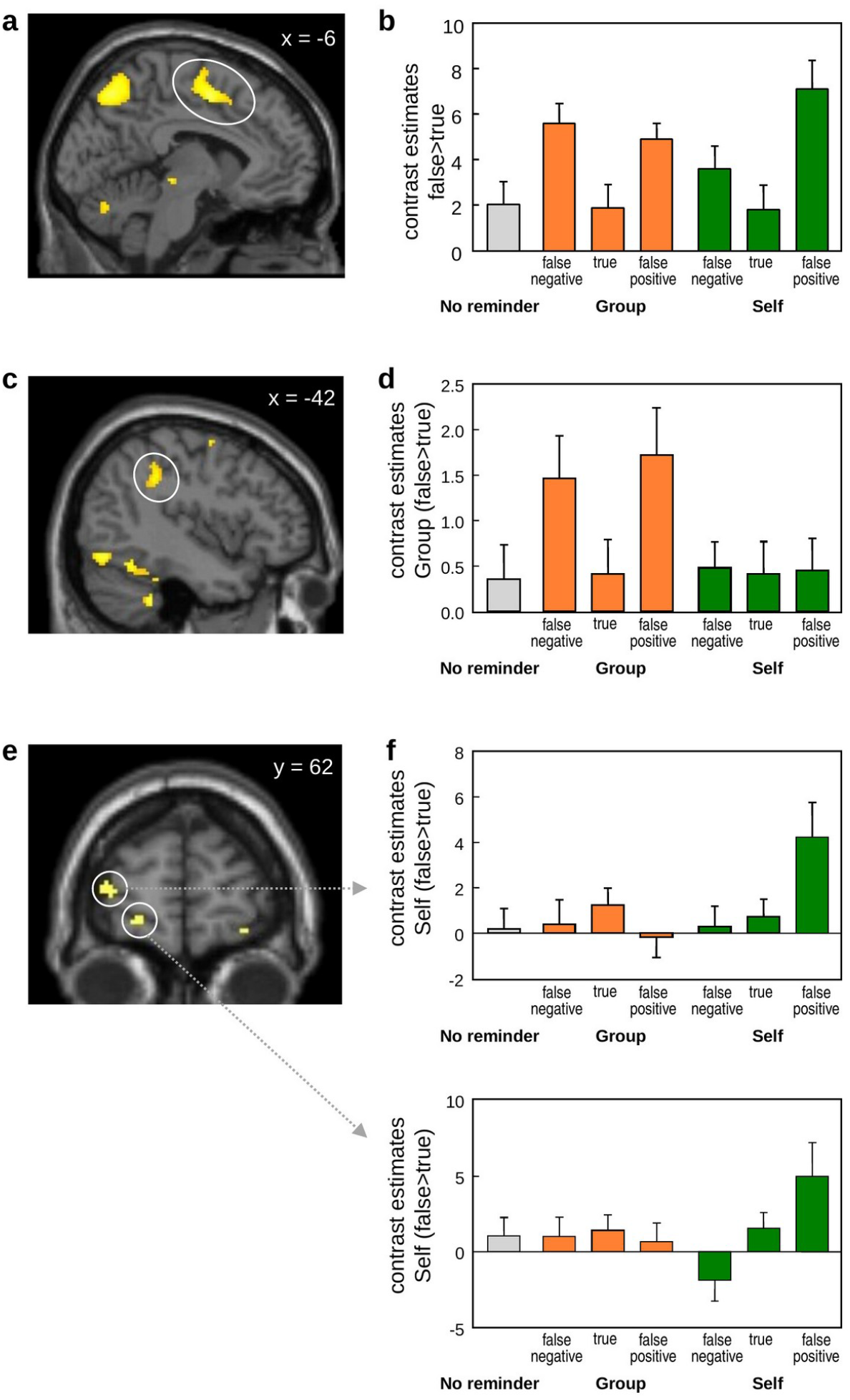
Conflict-related activation. (a,b) Regions more strongly activated for shifted than non-shifted reminders, common to the group and self conditions, including dmPFC (a; x/y/z coordinates = -6/0/50), with the corresponding effect sizes for false negative and false positive trials (b). (c,d) Regions showing conflict-related activity specific to the social condition, including left temporoparietal junction (c; -42/-40/36), and corresponding effect sizes (d). (e,f) Regions showing conflict-related activity specific to the self condition, including anterior vlPFC (e; -20/64/-6), and corresponding effect sizes (f). Statistical maps are displayed at the cluster-defining voxel-level threshold of *p < 0*.*001 (uncorrected); all mentioned clusters survive FWE cluster-level correction (p < 0*.*05;* Supplementary Table 1).

Next, we asked whether some regions preferentially reflected conflict for social or self information. We found conflict-like neural activity for social but not self conditions in the left temporoparietal junction (Fig. 3c; Supplementary Table 1). Again, activation increased similarly for both false negative and false positive trials (Fig. 3d). Conversely, anterior vlPFC (Fig. 3e; Supplementary Table 1) as well as inferior temporal lobe were more engaged in situations of conflict with individual previous ratings. Interestingly, anterior vlPFC showed increased activity only in the false positive trials but not in the false negative trials (Fig. 3f). Thus, some regions showed conflict-like activity particularly when social or individual information diverged from present participant judgment and even encoded the direction of this divergence.

#### Over-conformity and anti-conformity regardless of social and self condition

To identify the neural activations related to different responses to conflict, we modelled the data in a behavior-dependent fashion and compared parametric relations to the degree of behavioral adjustment for over-, under- and anti-conformity (GLM2). Irrespective of whether the conflict occurred from rating differences with the group or oneself, posterior bilateral vlPFC showed parametric activity increases with the degree of over-conformity (Fig. 4a,b; Supplementary Table 2). By contrast, activity in dmPFC (Fig. 4d,e; Supplementary Table 2) and left hippocampus (Fig. 4g,h; Supplementary Table 2) parametrically increased with the degree of anti-conformity. To investigate individual differences, we performed a brain–behavior correlation analysis, relating individual beta weights from the neural activity identified above to participant-specific degree of behavioral conformity types (average of group and self). The correlation plots show that the more strongly participants over-conformed, the more strongly the posterior vlPFC showed a parametric effect of over-conformity (Fig. 4c). On the other hand, the more participants anti-conformed, the stronger was the parametric effect of anti-conformity in dmPFC and hippocampus (Fig. 4f,i). Together, the extent to which participants increased the discrepancy between the current judgment and that of others or of the past self was associated with activation increase in dorsomedial regions of prefrontal cortex.

**Fig. 4.**
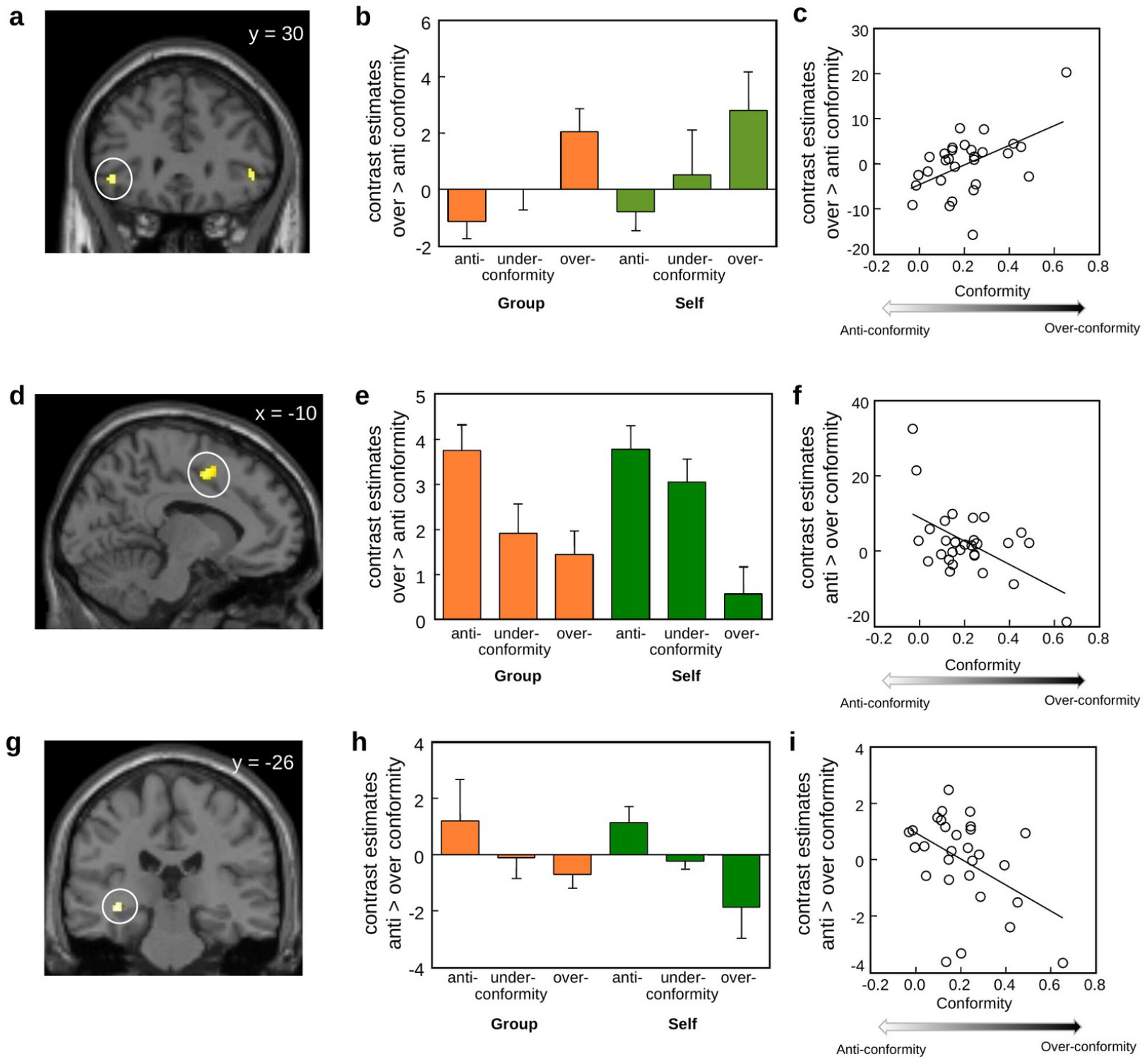
Activation related to over- and anti-conformity. (a–c) Posterior bilateral vlPFC (-46/30/-4) activity increasing parametrically with the degree of over-conformity (a,b) and the corresponding brain–behavior correlation (c). (d–f) dmPFC activity (-10/2/50) increasing with the degree of anti-conformity (d,e) and the corresponding correlation (f). (g–i) Left hippocampus activity (-36/-26/-6) increasing with anti-conformity (g,h) and the corresponding correlation (i). Statistical maps are displayed at the cluster-defining voxel-level threshold of *p < 0*.*001 (uncorrected); all mentioned clusters survive FWE cluster-level correction (p < 0*.*05; Supplementary Table 2)*.

#### Over-conformity and anti-conformity specific for social or self condition

Next, we searched for brain regions specific to group or self-conformity and found that right anterior vlPFC activity parametrically processed over-conformity when participants over-conformed to the group but not when they over-conformed to their own previous judgment (Fig. 5a,b; Supplementary Table 3). Finally, a left ventromedial prefrontal region was specific to over-conformity with oneself, while the left insula was specific to anti-conformity with oneself (Fig. 5d,e,g,h; Supplementary Table 3). Brain–behavior correlation plots show the individual tendencies for over- or anti-conformity specific to group or self (Fig. 5c,f,i). Thus, within the ventral prefrontal cortex, we found a dissociation such that more medial regions preferentially encoded the degree of conformity with oneself whereas more lateral regions preferentially encoded the degree of conformity with the group. Figure 6 summarizes these and all the other findings.

**Fig. 5.**
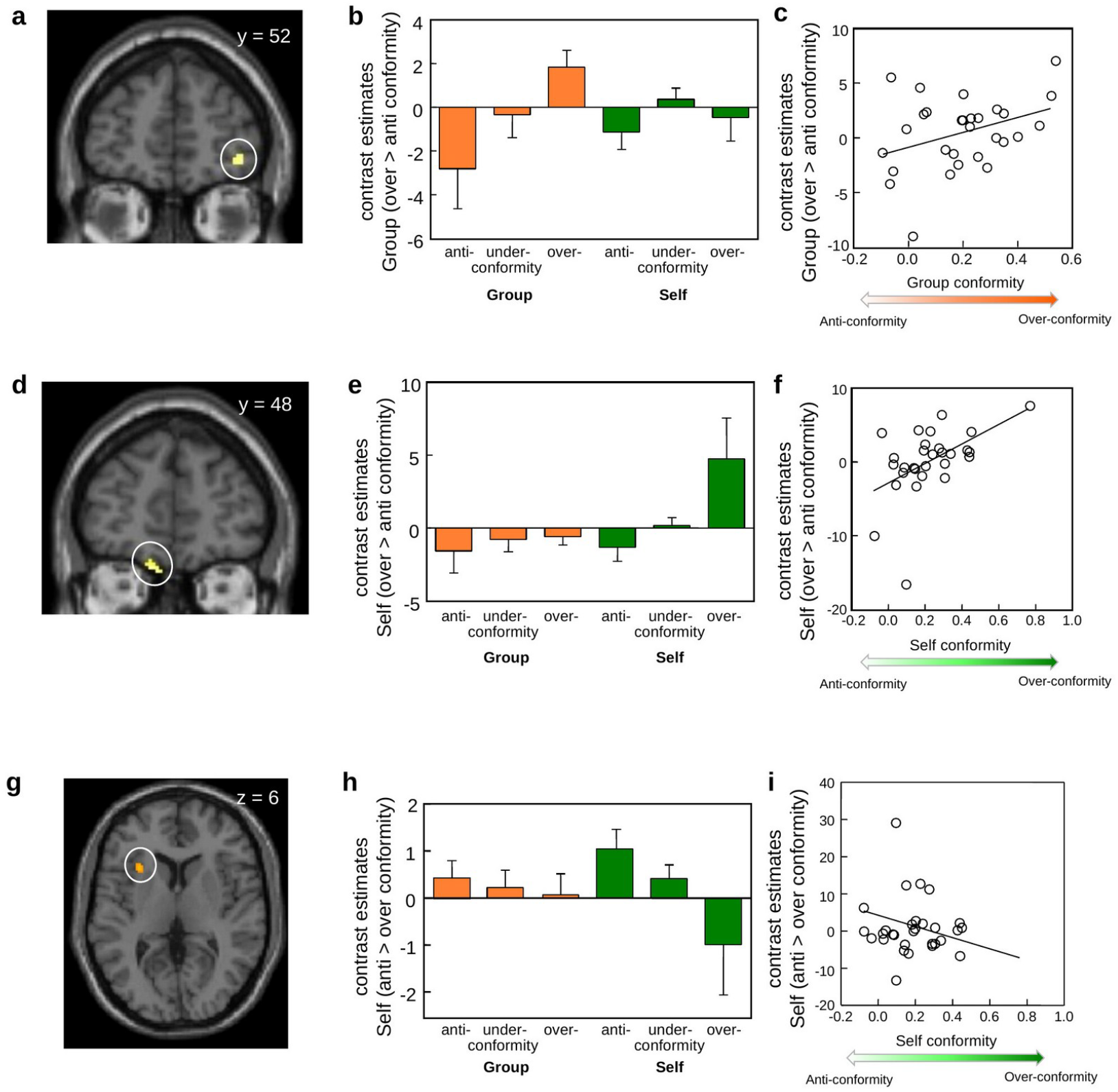
Group- and self-specific activation. (a–c) Right anterior vlPFC activity (40/52/-6) specific to over-conformity with the group (a,b) and the corresponding correlation (c). (d–f) Left ventromedial prefrontal cortex activity (-12/48/-24) specific to over-conformity with oneself (d,e) and the corresponding correlation (f). (g–i) Left insula activity (-30/22/6) specific to anti-conformity with oneself (g,h) and the corresponding correlation (i). Statistical maps are displayed at the cluster-defining voxel-level threshold of *p < 0*.*001 (uncorrected); all mentioned clusters survive cluster-level FWE-correction (p < 0*.*05; Supplementary Table 3)*.

**Fig. 6.**
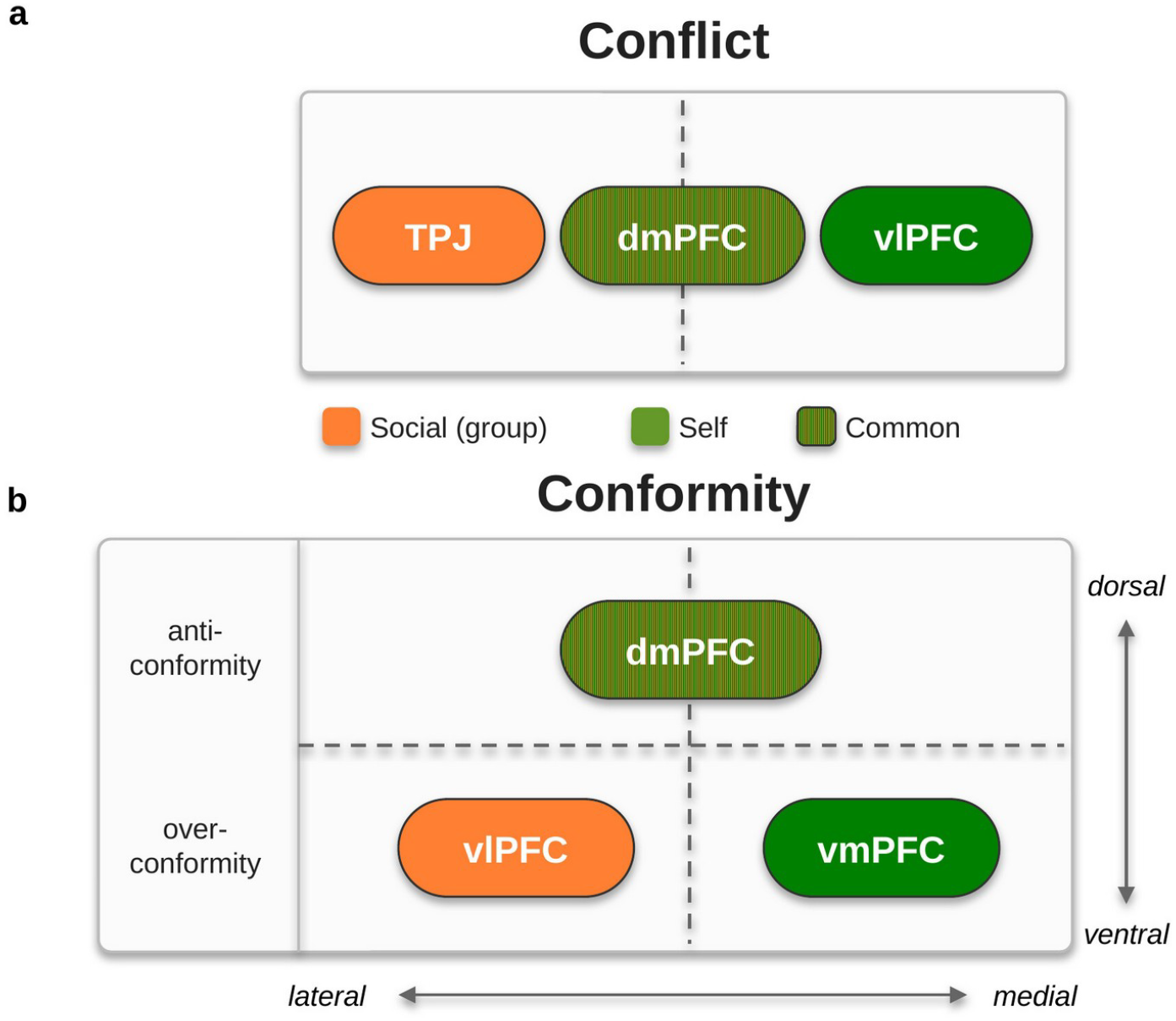
Schematic summary of main findings. Regions associated with conflict (a) and with conformity (b) across the social (group) and self conditions. The temporoparietal junction (TPJ) preferentially processed social conflict whereas ventrolateral prefrontal cortex (vlPFC) preferentially processed conflict with self. Dorsomedial prefrontal cortex (dmPFC) engaged in both forms of conflict. (b) Anti-conformity activated dmPFC, whereas over-conformity engaged vlPFC (social) and ventromedial prefrontal cortex (vmPFC; self). A possible interpretation of our findings is in terms of a medial–lateral (self versus social) and a dorsal–ventral (anti-versus over-conformity) organization within prefrontal cortex.

## Discussion

In this study, we investigated self-conformity and compared it with social conformity. Behaviorally, participants conformed to a similar degree to themselves and to the group; nonetheless, the brain partly dissociated the two forms of conformity. In particular, vmPFC (over-conformity with oneself), insula (anti-conformity with oneself), and vlPFC (over-conformity with the group) showed specific signals. By contrast, common parametric anti-conformity and conflict signals occurred in the dmPFC. Thus, while self-conformity shares neural architecture with social conformity, the brain also differentiates between the two.

In line with the notion that people desire to align not only with others^1–3^ but also with themselves, we found that participants showed self-conformity and that the propensity for self-conformity correlated with the propensity to conform to the group. Thus, in our task, participants changed their ratings of a face to a similar degree when either the social or their previous judgment deviated from their actual judgment. Conforming to the externally provided information reduces inconsistency with others and oneself, potentially increasing approval by others and oneself.

Similar to choice blindness^4^, participants failed to notice that the presented reminder about their previous judgment was incorrect and often conformed (or anti-conformed) to it. However, both self-conformity and social conformity were relatively short-lived as ratings returned to the initial ratings within two days after the main task. This finding is worth considering alongside our exploratory personality finding: conformity was stronger in participants with higher in agreeableness and conscientiousness (while low agreeableness was associated with anti-conformity). Thus, more stable personality traits may be predictive of the tendency to temporarily conform. Moreover, to the extent that conformity reflects a prosocial behavior, these findings converge with previous reports of correlations between agreeableness and prosocial behavior^14– 16^. Still, we note that these personality associations should be interpreted with caution given the modest sample size.

Pioneering studies on the neural mechanisms of social conformity highlight the important role of conflict-related regions and regions processing social rewards for adjusting to the opinion of others^3,9–13,18^. Consistent with these studies, we found conflict-like activation in dmPFC when false reminders were presented. Our findings add to this literature by revealing that these regions commonly encode conflict in the social and the self-condition.

The TPJ was more strongly engaged by conflicts with the group, while anterior vlPFC was more strongly engaged by discrepancy with oneself, at least when the reminder was more positive than the actual judgment. These findings add to reports of TPJ being involved in prosocial activity^19^ and encoding the prior choice of group members^20^ and of vlPFC being associated with conflict-driven cognitive control^21^. With regard to vlPFC, it is worth noting that its conflict-related processes are more pronounced in situations of self-relevance^22^. Taken together, our data suggest that TPJ and vlPFC are key regions for detecting conflict with others or oneself.

Different forms of conformity were associated with distinct neural activations. Compared to anti-conformity, over-conformity elicited stronger parametric activity increase in posterior vlPFC. In contrast, parametric anti-conformity more strongly related to dmPFC and hippocampus activity than parametric over-conformity. Converging evidence has associated vlPFC with susceptibility to social influence^23^. Indeed, greater activity in vlPFC correlates with participants’ willingness to alter their opinions when faced with discrepant attitudes of others^10^. On the other hand, dmPFC seems to relate to non-conformist behavior^24^ and individual differences in anti-social attitudes^25^. Together with our data, these findings suggest a dorsomedial-ventrolateral prefrontal axis that orchestrates the individual tendency to accommodate or antagonize others.

While medial parts of medial prefrontal cortex related more to over-conformity with oneself, the insula preferentially related to anti-conformity with oneself. The vmPFC and anterior insula have both been associated more strongly with self-related than other-related judgments^26,27^. Moreover, they were also respectively implicated in serving self-reference functions in the default mode network^28^ or the salience network^29^. Our data may suggest that it could be worth assessing whether anti-conforming with oneself is more salient than over-conforming with oneself.

Taken together, our results suggest that medial versus lateral parts of ventral prefrontal cortex differentially relate to the attribution of judgments to self or others, whereas dorsomedial versus ventrolateral prefrontal cortex relates to the tendency to accommodate or antagonize others. These dissociations characterize prefrontal cortex with a multifaceted role in determining the target and degree of conformity.

## Methods

### Participants

We recruited 29 healthy male participants (mean age: 31.0 years; range: 20–47 years) from the Fukushima Medical University community. All participants were right-handed with no history of neurological, psychiatric, or auditory dysfunction. They all had normal or corrected to normal vision. The study was conducted in accordance with the Declaration of Helsinki and approved by the Research Ethics Committee of the Fukushima Medical University (approved No. 2056). Written informed consent was obtained from each participant. In Fig. 1 the face stimuli are shown as schematic placeholder icons rather than the actual photographs, which depicted real faces (see Stimuli). All participants were paid a fixed total of JPY 6,000 (USD 55) upon completion of the experiment.

### Experimental design

#### Stimuli

We used 360 digital photos of Japanese female faces from free internet sources (https://woman.excite.co.jp/, accessed in 2015, by permission to use only in this study). All images were modified from the original (i.e., face-cropped, background-removed and resized). Stimulus presentation and timing was implemented using Cogent 2000 (Wellcome Department of Imaging Neuroscience, London, United Kingdom) and Matlab 7 (Mathworks). The images were presented on an MR-compatible LCD monitor and viewed through a mirror mounted on the head coil. Participants’ responses were obtained with a MR-compatible trackball.

#### Task and procedure

Participants completed a simple face attractiveness rating task in the fMRI scanner on three different occasions. On the first day in the scanner (pre-rating task; Fig. 1a), they were asked to rate the 360 photos for facial attractiveness one by one on a visual analog scale (VAS) using a trackball. The initial position of the cursor was random. The extremes of the VAS were labeled Low and High, and the position of the labels was flipped for half of the participants. Responses had to occur within 5000 ms and intertrial intervals ranged between 500 and 1500 ms. The pre-rating task consisted of two scanning sessions (180 stimuli each) and lasted about 36 min in total.

The second day in the scanner (main rating task; Fig. 1b) took place one week later. Participants rated 252 faces used in the pre-rating task. The main rating task comprised three trial types. First, faces were accompanied by the average rating of the social group, labeled “Average”. This information ostensibly reflected how attractive a previous group of 28 participants had found that face. Second, faces appeared together with the ostensible previous rating of the participant, labeled as “You”. Third, faces were presented on their own without a reminder (as in the pre-rating task), labeled with “No reminder”. The “Average” and “You” ratings were determined based on each participant’s pre-rating. For both the “Average” and “You” trials, the reminder was derived from the participant’s pre-rating: in one third it was the actual pre-rating (true trial), and in the remaining two thirds it was shifted below the participant’s pre-rating (“false negative”) or above it (“false positive”; for more details on the shifting procedure, please see supplementary information, Supplementary Fig. 1a,b). For “Average” trials, this reminder was presented as the group (“Average”) rating. Social conformity and self-conformity were measured as change in ratings from pre-rating to main rating. The main rating task lasted about 26 minutes.

Finally, participants came to the scanner a third time to perform the post-rating task (Fig. 1c). Six participants were scanned two days after the main task and the remaining 23 participants performed the task one week after the main task. As there was no difference between participants who performed the post-rating task two or seven days after the main rating task, we combined all the data. As in the pre-rating task, participants rated the original 360 faces without reminders. Comparing ratings across tasks allowed us to assess the temporal stability of conformity. Participants next answered questions regarding the study. None of them reported that they realized or even suspected a shift of the reminder ratings displayed during the main task. Finally, they completed the revised NEO Personality Inventory (NEO-PI-R).

### Data acquisition and preprocessing

#### Data acquisition

We acquired all neuroimaging data on a 3-T Siemens Biograph mMR scanner (Siemens, Erlangen, Germany) with a 12-channel head matrix coil at Fukushima Medical University. Functional images were acquired using a gradient-echo echo-planar imaging (EPI) sequence (TR = 3000 ms; TE = 30 ms; FA = 90°; FOV = 192 mm; 64 × 64 matrix; 45 slices; slice thickness = 3.0 mm, voxel size = 3.0 × 3.0 × 3.0 mm). The pre- and post-rating tasks each consisted of 371 functional volumes, whereas the main rating task consisted of 520 functional volumes. After completion of the functional runs, a high-resolution structural T1-weighted image was acquired (TR = 1800 ms, TE = 1.99 ms, FA = 90°; FOV = 250 mm; 256 × 256 matrix; 176 slices; slice thickness = 1 mm, voxel size = 1.0 × 1.0 × 1.0 mm).

#### Pre-processing procedures

Preprocessing and data analysis were performed using Statistical Parametric Mapping 12 (SPM12) software. The first 3 functional scans of each session were discarded to allow for magnetic saturation. CONN served to detect outliers (global signal z value threshold = 5 SDs, subject-motion threshold = 0.9 mm) and determine global gray matter signal. Functional images were realigned, unwarped, and slice-time corrected. We checked for motion artifacts using the ART-based scrubbing method^17^. Gray matter, white matter, and cerebrospinal fluid were segmented, and the functional data were normalized to the Montreal Neurological Institute template. Finally, the data were spatially smoothed with a Gaussian kernel set at 8 mm full width at half-maximum.

### fMRI data analysis

#### Identification of conflict regions (GLM1)

First, we aimed to identify brain activation related to the detection and monitoring of conflict between participants’ current ratings and false information in the main task. We therefore performed event-related general linear model (GLM) analyses at the time of stimulus presentation. Seven regressors corresponding to the different trial types (true, false negative or false positive trials in both “Average” and “You”; and “No reminder”) were modeled as stick functions (duration of 0 s). Participant-specific motion parameters were also modeled as covariates of no interest. A high-pass filter with a cut-off period of 128 s was used to remove low-frequency drifts. We performed two-way ANOVA in the second level analysis on contrasts between false and true trials. Regions specific for conflicts with social or self-ratings were identified separately. Clusters were formed with a voxel-level threshold of *p < 0*.*001 (uncorrected) and are reported together with their whole-brain cluster-level family-wise error (FWE)-corrected p*-values (Supplementary Table 1). Only clusters surviving whole-brain FWE cluster-level correction (*p < 0*.*05) are described in the main text*.

#### Different conformity types (GLM2)

To identify brain activation related to subsequent change in behavior, we used the brain activation from the onset of the stimulus until rating was completed because we expected the neural effects of conformity to occur during the rating response (variable duration, average = 3.5 s). According to the direction of behavioral conformity (change in ratings from pre-rating to main rating), we grouped trials into three types. First, trials where participants rated a given face even more extremely than the indicated rating of the group or themselves (over- conformity); second, trials with a rating that did not exceed the indicated rating (under-conformity); and third, trials in which participants rated the face in the opposite direction of the indicated rating (anti-conformity). Accordingly, GLM2 comprised 15 regressors in total: over-/under-/anti-conformity for false negative or false positive trials in both “Average” and “You” conditions, true trials in both “Average” and “You”, and “No reminder” conditions. We used the degree of conformity as parametric modulator for each regressor where conformity was possible. This allowed us to detect linear activation changes related to trial-by-trial degree of conformity. Six motion parameters were included as covariates of no interest. We identified group conformity and self-conformity as well as over- or anti-conformity specific regions separately on the second level and performed two-way ANOVA contrasting over-conformity against anti-conformity for both group and self conditions. As in GLM1, clusters were formed with a voxel-level threshold of *p < 0*.*001 (uncorrected) and are reported with their whole-brain cluster-level FWE-corrected p*-values (Tables S2 and S3). Only clusters surviving whole-brain FWE cluster-level correction (*p < 0*.*05) are described in the main text*.

## Acknowledgements

This work was supported by the Grant-in-Aid for JSPS Fellows 12J03478 and JSPS KAKENHI Grant Number 19K07803. PNT was supported by the Swiss National Science Foundation (grants 10.001.677 and 207613). We thank Hitoshi Kubo for assistance with scanning.

## Author contributions statement

J.F. conceived and designed the study, performed the experiments and the analyses, and wrote the paper with P.N.T. All authors contributed to the interpretation of the results and reviewed and approved the manuscript.

## Additional information

Supplementary Information (Supplementary Figs. 1–2 and Supplementary Tables 1–3) is available as a separate file accompanying this preprint.

Competing Interests: The authors declare no competing interests.

## Data availability

De-identified behavioural and personality data that support the findings of this study are available at the Open Science Framework [https://osf.io/8kfcn]. The group-level (unthresholded) statistical maps will be made publicly available on NeuroVault upon publication and are available from the corresponding author on reasonable request in the interim. The individual-level raw neuroimaging data and the facial stimuli are not publicly available because they contain information that could compromise participant privacy and because the stimuli were licensed for use in this study only; they are available from the corresponding author upon reasonable request.

## Code availability

The custom analysis code used in this study (implemented in SPM12 and MATLAB) is publicly available at GitHub [https://github.com/jurifujiwara/self-conformity-fmri] and archived at Zenodo [https://doi.org/10.5281/zenodo.21333864].

