## Supplementary Information for "Neural mechanisms of self-conformity"

†Deceased.

**Supplementary information**

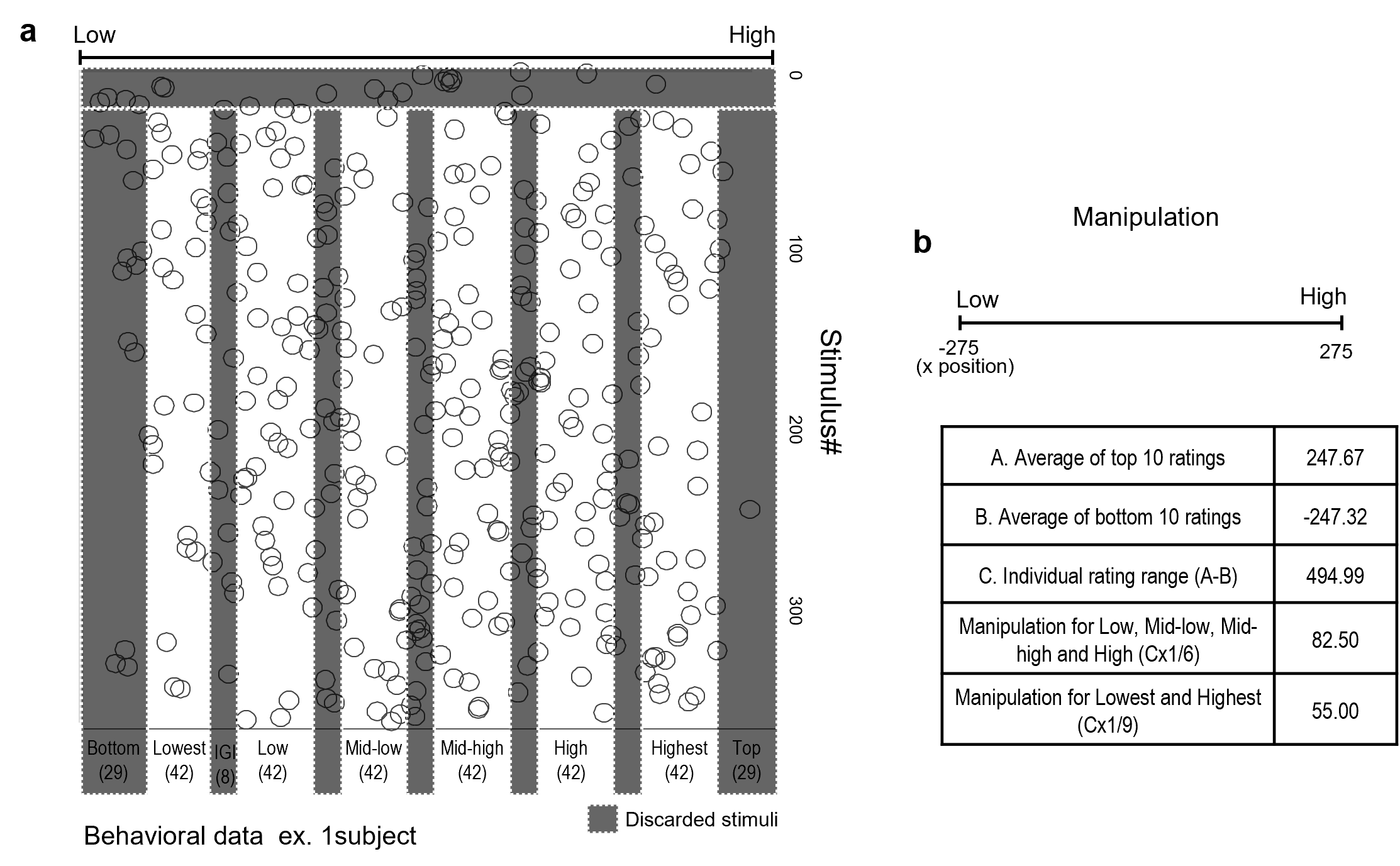

**Supplementary Fig. 1.**

**Stimulus grouping.** On the basis of their individual pre-ratings, we systematically split all stimuli into six groups (42 stimuli for each group: Lowest, Low, Mid-low, Mid-high, High and Highest. Between each pair of adjacent groups, eight stimuli were discarded to separate them). Stimuli from these groups were presented pseudo-randomly in the main rating task. Before grouping, we discarded the 10 stimuli participants rated first and the extremely high and low rated stimuli (29 stimuli from top and 29 stimuli from bottom) to allow for initial rating adjustment and prevent ceiling or flooring issues for the manipulation used in the main rating task. We used the remaining 252 stimuli in the main experiment (Supplementary Fig. 1a).

**Task manipulation.** Both social and individual trials consisted of two conditions; true (reminder same as previous rating; 33 % of the trials) or false (reminder shifted above or below their previous rating; 33 % of the trials each). False ratings were manipulated using the following criteria: participant’s previous ratings were pseudorandomly shifted above or below constrained by individual rating range which was calculated as follows (average of top 10 ratings – average of bottom 10 ratings) x *1/6 in non-extreme group or *1/9 in lowest and highest groups (Supplementary Fig. 1b). False ratings for group trials were manipulated exactly as individual false ratings but described as “Average” group ratings.

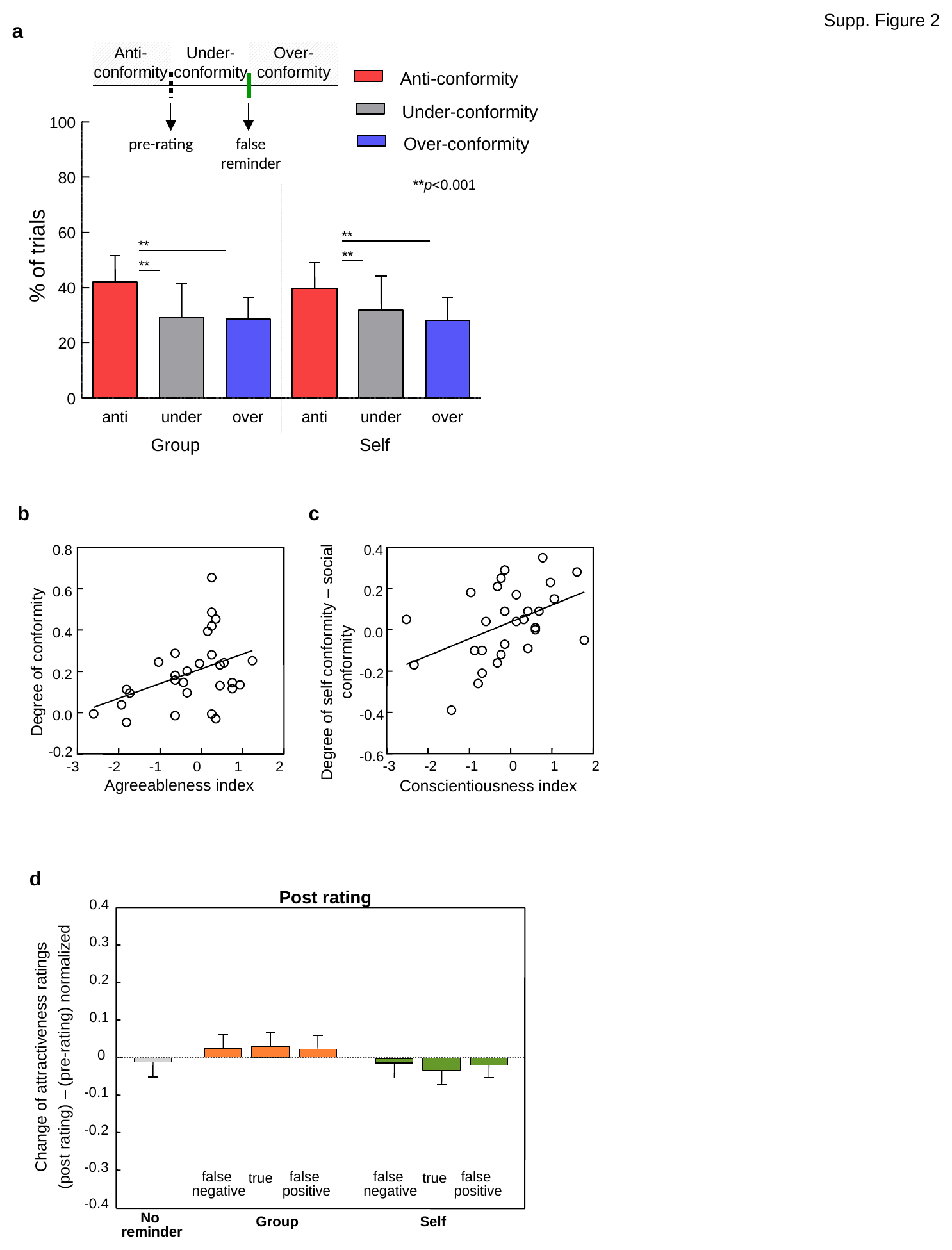

**Supplementary Fig. 2.** Temporal stability and distribution of conformity. (a) Frequency of over-, under- and anti-conformity across participants (F(2,56)=10.17, p < 0.0005). (b) Relationships between overall conformity and agreeableness and (c) between self-relative to group conformity and conscientiousness. (d) Conformity measured in the post-rating task compared with the main rating task, showing a return towards the initial ratings within a few days (compare with Fig. 2a of main manuscript).

### Supplementary Table 1. Conflict-related regions

| Brain region | L/R | x | | y | | z | Peak Z | | Cluster p(FWE) | k (voxels) |
| --- | --- | --- | --- | --- | --- | --- | --- | --- | --- | --- |
| **Fig. 3a, Conflict common region** | | | | | | | | | | |
| Dorsomedial prefrontal cortex | L | -6 | 0 | | 50 | | | 7.51 | <0.001 | 2545 |
| Dorsolateral prefrontal cortex | L | -40 | 6 | | 30 | | | 5.58 | <0.001 | 293 |
| Dorsolateral prefrontal cortex | R | 50 | 8 | | 26 | | | 5.52 | <0.001 | 135 |
| Orbitofrontal cortex | R | 26 | 56 | | -16 | | | 5.09 | 0.009 | 15 |
| Midbrain | R | 4 | -26 | | -10 | | | 5.58 | 0.001 | 54 |
| Precuneus | L | -20 | -68 | | 50 | | | 7.61 | <0.001 | 1545 |
|  | L | -24 | -62 | | 40 | | | 7.30 |  |  |
| Precuneus | R | 14 | -64 | | 44 | | | 7.03 | <0.001 | 1690 |
| Superior parietal lobule | R | 32 | -54 | | 54 | | | 6.79 |  |  |
| Cerebellum | L/R | 0 | -56 | | -34 | | | 6.18 | <0.001 | 81 |
| **Fig. 3c, Conflict group-specific region** | | | | | | | | | | |
| Superior frontal gyrus | L | -20 | -2 | | 56 | | | 5.78 | <0.001 | 849 |
| Superior frontal gyrus | R | 26 | 4 | | 56 | | | 5.75 | <0.001 | 759 |
| Dorsomedial prefrontal cortex | L | -6 | 2 | | 50 | | | 4.48 | <0.001 | 83 |
| Precuneus | L | -14 | -62 | | 60 | | | 5.61 | <0.001 | 1626 |
| Supramarginal gyrus (TPJ) | L | -42 | -40 | | 36 | | | 3.58 |  |  |
| Precuneus | R | 14 | -62 | | 46 | | | 5.09 | <0.001 | 833 |
| Occipital gyrus | L | -38 | -70 | | -14 | | | 4.72 | <0.001 | 442 |
| Occipital gyrus | R | 40 | -76 | | -14 | | | 5.02 | <0.001 | 419 |
| **Fig. 3e, Conflict self-specific region** | | | | | | | | | | |
| Superior frontal gyrus | R | 28 | 0 | | 56 | | | 4.28 | 0.129 | 45 |
| Ventrolateral prefrontal cortex | L | -20 | 64 | | -6 | | | 4.45 | 0.015 | 109 |
| Ventrolateral prefrontal cortex | R | 38 | 42 | | -6 | | | 4.12 | 0.032 | 63 |
| Inferior temporal lobule | R | 46 | -54 | | -14 | | | 4.45 | 0.008 | 153 |
| Precuneus | L | -22 | -62 | | 38 | | | 4.89 | <0.001 | 875 |
| Precuneus | R | 14 | -66 | | 42 | | | 4.95 | <0.001 | 705 |
| Occipital gyrus | L | -34 | -82 | | -2 | | | 4.79 | <0.001 | 400 |
| Cerebellum | L/R | 0 | -56 | | -34 | | | 4.55 | 0.178 | 35 |

Regions are reported at a cluster-defining voxel-level threshold of *p* < 0.001 (uncorrected); the Cluster *p*(FWE) column gives the whole-brain cluster-level family-wise error (FWE)-corrected value. Clusters surviving whole-brain FWE correction (*p* < 0.05) are those described in the main text.

### Supplementary Table 2. Conformity-related regions

| Brain region | L/R | x | y | z | Peak Z | Cluster p(FWE) | k (voxels) |
| --- | --- | --- | --- | --- | --- | --- | --- |
| **Fig. 4a, Over-conformity common region** | | | | | | | |
| Ventrolateral prefrontal cortex | L | -46 | 30 | -4 | 4.50 | 0.003 | 48 |
| Ventrolateral prefrontal cortex | R | 48 | 28 | -2 | 4.35 | 0.010 | 41 |
| Anterior cingulate gyrus | L | -4 | 20 | 20 | 3.96 | 0.052 | 18 |
| Fusiform gyrus | R | 30 | -16 | -34 | 4.10 | 0.002 | 68 |
| **Fig. 4d,g, Anti-conformity common region** | | | | | | | |
| Superior frontal gyrus | R | 26 | -6 | 46 | 4.69 | 0.022 | 122 |
| Dorsomedial prefrontal cortex | L | -10 | 2 | 50 | 5.51 | 0.008 | 165 |
| Hippocampus | L | -36 | -26 | -6 | 4.22 | 0.040 | 28 |
| Precuneus | L | -20 | -62 | 40 | 4.51 | 0.016 | 164 |
| Precuneus | R | 20 | -60 | 46 | 4.27 | 0.024 | 107 |
| Occipital gyrus | L | -30 | -84 | -8 | 4.30 | 0.001 | 206 |
| Occipital gyrus | R | 36 | -78 | -12 | 4.16 | 0.031 | 31 |

### Supplementary Table 3. Condition-specific regions

| Brain region | L/R | x | y | z | Peak Z | Cluster p(FWE) | k (voxels) |
| --- | --- | --- | --- | --- | --- | --- | --- |
| **Fig. 5a, Over-conformity group-specific region** | | | | | | | |
| Ventrolateral prefrontal cortex | R | 40 | 52 | -6 | 4.42 | 0.010 | 22 |
| Ventrolateral prefrontal cortex | L | -44 | 34 | -4 | 4.26 | 0.032 | 15 |
| Inferior temporal gyrus | R | 30 | 4 | -44 | 4.30 | 0.010 | 39 |
| Occipital gyrus | R | 28 | -78 | 22 | 4.39 | 0.004 | 66 |
| **Fig. 5d, Over-conformity self-specific region** | | | | | | | |
| Ventromedial prefrontal cortex | L | -12 | 48 | -24 | 4.22 | 0.042 | 17 |
| Precentral gyrus | L | -18 | -10 | 70 | 4.52 | 0.025 | 48 |
| Supplementary motor area | R | 8 | 16 | 62 | 5.02 | 0.012 | 60 |
| **Fig. 5g, Anti-conformity self-specific region** | | | | | | | |
| Dorsomedial prefrontal cortex | L | -4 | -2 | 48 | 6.80 | 0.004 | 96 |
| Middle cingulate gyrus | L | -8 | -30 | 32 | 3.96 | 0.113 | 15 |
| Insula | L | -30 | 22 | 6 | 4.24 | 0.006 | 22 |

Regions are reported at a cluster-defining voxel-level threshold of *p* < 0.001 (uncorrected); the Cluster *p*(FWE) column gives the whole-brain cluster-level family-wise error (FWE)-corrected value. Clusters surviving whole-brain FWE correction (*p* < 0.05) are those described in the main text.
